# New insights on species divergence in red panda

**DOI:** 10.1101/2020.08.27.268607

**Authors:** Bheem Dutt Joshi, Supriyo Dalui, Sujeet Kumar Singh, Tanoy Mukherjee, Kailash Chandra, Lalit Kumar Sharma, Mukesh Thakur

## Abstract

With the recent classification of red panda into two phylogenetic species, we propose ‘*Siang river*’ as a potential boundary for species divergence between the Himalayan red panda (*Ailurus fulgens*) and the Chinese red panda (*Ailurus styani*). Bayesian based phylogeny and MJ network splited all the sequences into two distinct clusters in accordance to the origin of the samples collected from the east and west side of the Siang river. The clade 1, that represented Himalayan red panda, was formed by inclusion of samples originated from the north West Bengal, Sikkim, and central Arunachal Pradesh and South Tibet. While, clade 2 represented Chinese red panda by inclusion of the samples originated from Dibang valley of eastern Arunachal Pradesh, India and Southwest China. We suggest being associated with diverse habitats, threats and transboundary distribution, both species of red panda require regional as well as multilateral cooperation for making species survival plan.

## Introduction

Among one of most elusive and endangered species-red panda, recently classified into two species, *i.e*. Himalayan red panda (HRP; *Ailurus fulgens*) and Chinese red panda (CRP; *Ailurus styani*)^1^. Of which, HRP distribution is reported in all along the Himalayan region of Nepal, North West Bengal (Darjeeling and Kalimpong districts), Sikkim and Arunachal Pradesh States in India, Bhutan, Northern Myanmar, Tibet and Western Yunnan^2^. Whereas the CRP panda distributed in the Yunnan and Sichuan provinces of China^1^ and Nujiang river was proposed as a geographic boundary between the two species^3^. Unfortunately, both the species of red panda have rapidly lost suitable habitat across the range due to landscape fragmentation, change in the land use patterns, anthropogenic activities^4–7^. Also red panda has been facing poaching threats to supply the demands for skins, parts and for traditional medicine^6,8^. As a conservation measure, red panda is categorized as ‘*Endangered*’ under IUCN Red list and listed under *Schedule I* of the Wildlife (Protection) Act, 1972 of India. Red panda mostly inhibits in the temperate, montane forest (i.e. oak mixed, mixed broad-leaf conifer and conifer) and typically an obligatory bamboo feeder^4,9^. Recently, Hu et al. (2020) proposed Yalu Zangbu river that originates at Angsi Glacier in the western Tibet, southeast of Mount Kailash and Lake Manasarovar, is the geographic barrier for species divergence in red panda^1^. However, these authors did not include red panda samples from India and made inferences to be validated by incorporating samples from India and Bhutan.

Since, delineate species boundaries at fine scale is imperative to support species oriented conservation plans and to avoid inappropriate conservation measures, e.g. a misleading detrimental interbreeding between two distinct species in captivity or translocation of individuals in one another species range^10,11^. We aimed this study to investigate the species boundary between the two phylogenetic species of red panda by incorporating samples collected from the various extent of distribution in the Indian Himalayan Region (IHR).

## Methods

We collected 132 fecal samples that included 29 samples from the North West Bengal, 28 from Sikkim in the Central Himalaya and 75 from the Arunachal Pradesh in the Eastern Himalayas. Genomic DNA was extracted using QIAamp Fast DNA Stool Mini Kit (QIAGEN Germany) following manufacturer’s instructions. We sequenced 440 bp control region fragment using red panda specific primers^14,15^. PCR amplification reaction performed on Veriti thermal cycler (Applied Biosystems, USA) with total volume of 10 μl comprising of 20-30 ng template DNA, 1U Taq polymerase (Takara), 10X PCR buffer, 1 mM MgCl_2_, 2.5 mM dNTPs mix, 0.1 μM of each primer, 0.1 μg/μl BSA. Thermal cycling conditions were as follows: an initial denaturation step at 94°C for 5 min, followed by 35 cycles at 94°C for 30s, 50°C for 45s and 72°C for 1 min. The final extension was at 72°C for 10 min. Sequencing was performed using Big-Dye Terminator Cycle Sequencing Kit v.3.1 (Thermo Scientific, USA) on ABI 3730 Genetic analyzer (Applied Biosystems, USA).

We downloaded 44 complementary sequences of red panda available in the public domain (Table S2). We aligned all sequences using Clustal W implemented in BioEdit^16^ and intra-specific genetic distances were estimated using MEGA X^17^. We chose the best suitable nucleotide substitution models based on the Akaike information criterion (AIC)^18^ and the Bayesian information criterion (BIC) using MrModeltest v2.3^19^. Bayesian based phylogeny was 20 performed following Hasegawa-Kishino-Yano (HKY) mutation model in BEAST 2 considering racoon (*Procyon lotor; GenBank ID*: AB291073.1) as an outgroup. The phylogenetic analysis was performed for 200,000,000 generations each, sampling every 10,000 generations and performance was checked using Tracer v1.5. The resulting trees were combined using Log Combiner, and the consensus tree was generated using TreeAnnotator, after a burn-in of 25% maximum clade credibility (MCC) tree^20^. The MCC tree was visualized in the FigTree v.1.3.1^21^.

## Results and discussion

Altogether, 88 sequences of which 44 generated in the present study that covered Central (Darjeeling in North West Bengal and Sikkim) and Eastern Himalayas (Arunachal Pradesh), yielded 53 haplotypes (18 novel haplotypes generated from IHR, and are submitted to GenBank; MT891293-MT891310) with sequence divergence ranging from 0.005-0.031. The Bayesian phylogeny clustered all the sequences into two major clades, clade 1 and 2. Interestingly, all the samples collected from the West of Siang river, including samples originated from Central Himalayas (Darjeeling in North West Bengal, Sikkim and South Tibet) and Western & Central Arunachal Pradesh in the Eastern Himalayas assigned to the clade 1 while six samples that contributed four haplotypes and collected from the Dibang valley clustered with the sequences of the CRP in clade 2 (Fig. 1 a and b). Noteworthy, samples collected from Dibang valley were actually located on the east of Siang river, which indicated that Siang river has been the potential barrier of species divergence in red panda while, clade 1 and clade 2 correspond to the HRP and CRP, respectively (Fig. 1a and b). The median-joining network also spanned with two groups, one for HRP and another for CRP which was in coherence to the Bayesian phylogeny. Here also, the four haplotypes of Dibang valley originated samples clustered along with the CRP (Fig. 1a; Dibang valley originated haplotypes shown in red color). Earlier, Wei et al. proposed that Nujiang river is a geographic boundary for the two species of red panda^3^ but, Hu et al. recently suggested that Yalu Zangbu river, which is the longest and the widest river of Tibet region, is a potential barrier which diverged HRP and CRP^1^. It is noteworthy that Yalu Zangbu is an upstream river to Siang river which originates at Angsi Glacier in western Tibet, southeast of Mount Kailash and Lake Maanasarovar, and later feeds to the Siang river in the state of Arunachal Pradesh, India. Hu et al. study left a substantial geographical gap in sampling the HRP. The present study by inclusion of samples collected from IHR and availing competent data from public domain, demonstrated that Siang river which flows in the downstream from the Yalu Zangbu river and located upstream to Brahmaputra river, is the actual potential barrier for species divergence in red panda (Fig. 1c). Further, one individual sampled on the edge of west of Siang river and clustered with HRP clade, exhibited high sequence divergence (0.016-0.031) from the samples collected from Dibang valley (Table S1). This evident to support for occurring two phylogenetic species of red panda in India, with Siang river in Arunachal Pradesh as a potential geographic barrier. Since, both the species encompasses transboundary distribution, where HRP distributed in Nepal, India, Bhutan and Tibet and the CRP distributed in India (East of Siang river in Arunachal Pradesh), Southwest China and Myanmar^1^. Both species need attention at regional level to explore the possible suitable habitats of red panda all along the east-west side of Siang river and we appeal to formulate species oriented transboundary conservation action plans. Also from the potential ancient refugia that have considered northeast hill of IHR as biodiversity hotspot ^12,13^, extensive surveys are required to capture such speciation events in other existing endangered species that will aid in timely adopting appropriate conservation strategies for the threatened and elusive taxon.

**Figure 1.**
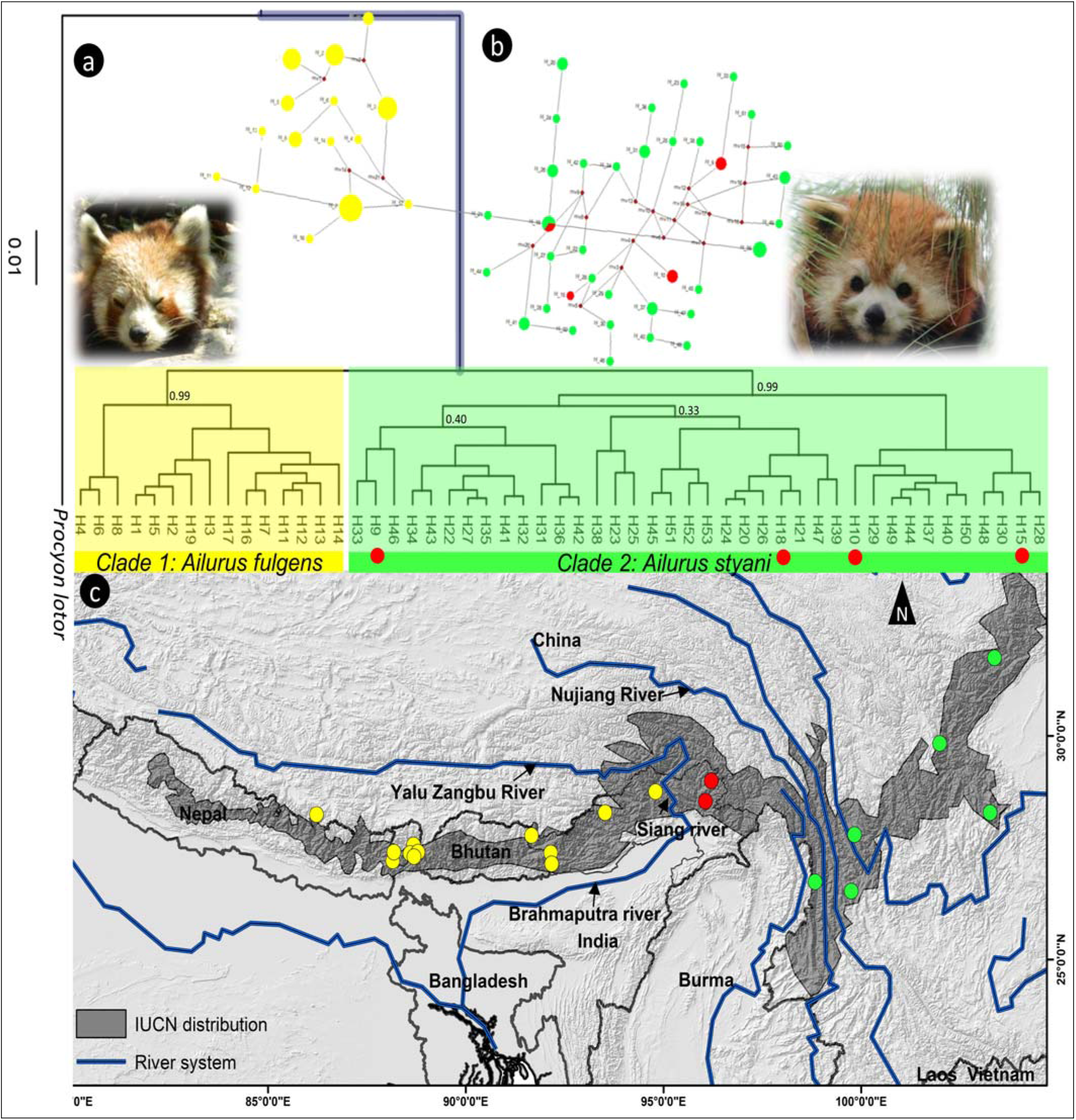
Depiction of red panda species divergence covering samples from the entire distribution range. a) Phylogenetic relationship of two red panda species in the central and eastern Himalaya with posterior probability depicted above the clade (yellow shaded clade 1 represented cluster of Himalayan red panda samples while green shaded clade 2 represented the cluster of Chinese red panda samples). Four samples collected in the east of Siang river but identified as Chinese red panda are highlighted in red dots. b). Median-joining haplotype network tree that correspond to the Bayesian phylogeny. The overlaid photographs are Himalayan red panda (left side) and Chinese red panda (right side) taken by the authors during field surveys. c). map showing the geographical distribution of red panda samples used in the present study on the overlaid IUCN defined range. The yellow color dots represented locality of Himalayan red panda specific haplotypes found in the samples collected on the west of Siang river while the red color dots represented haplotypes of Chinese red panda collected in the east of Siang river.

## Acknowledgements

Authors thank Principal Chief Conservator of Forest (PCCF) and Chief Wildlife Warden (CWLW) of red panda range States in India (Arunachal Pradesh, Sikkim and West Bengal) for granting necessary permission for undertaking field surveys and data collection. The study was supported by the DST - INSPIRE FACULTY SCHEME (Grant No. DST/INSPIRE/04/2016/002246) awarded to Dr Mukesh Thakur and funds received by the NMHS-Large Grant project of the Ministry of Environment, Forest and Climate Change, New Delhi (Grant No. NMHS/2017-18/LG09/02/476).

