## Supplementary files for "New insights on species divergence in red panda"

Table S1. Sequence divergence between the samples collected from the West Siang and Dibang valley.

|  | West Siang | Dibang-1 | Dibang-2 | Dibang-3 | Dibang-4 |
| --- | --- | --- | --- | --- | --- |
| West Siang |  |  |  |  |  |
| Dibang-1 | 0.031 |  |  |  |  |
| Dibang-2 | 0.031 | 0.023 |  |  |  |
| Dibang-3 | 0.031 | 0.023 | 0.009 |  |  |
| Dibang-4 | 0.016 | 0.016 | 0.016 | 0.021 |  |

Table S2. Samples details of Chinese Red Panda and Himalayan Red Panda

| **Haplotype** | **Accession NO** | **Locality** | **State** | **Country** | **Reference** | **Species ID** |
| --- | --- | --- | --- | --- | --- | --- |
| H1 | MT891293 | Darjeeling zoo | West Bengal | India | Present study | *A.fulgens* |
| H2 | MT891294 | West Sikkim | Sikkim | India | Present study | *A.fulgens* |
| H2 | MT891294 | Singalila | West Bengal | India | Present study | *A.fulgens* |
| H2 | MT891294 | Singalila | West Bengal | India | Present study | *A.fulgens* |
| H2 | MT891294 | East Sikkim | Sikkim | India | Present study | *A.fulgens* |
| H2 | MT891294 | Neora Valley NP | West Bengal | India | Present study | *A.fulgens* |
| H2 | MT891294 | Neora Valley NP | West Bengal | India | Present study | *A.fulgens* |
| H3 | MT891295 | West Sikkim | Sikkim | India | Present study | *A.fulgens* |
| H3 | MT891295 | East Sikkim | Sikkim | India | Present study | *A.fulgens* |
| H3 | MT891295 | West Sikkim | Sikkim | India | Present study | *A.fulgens* |
| H3 | MT891295 | East Sikkim | Sikkim | India | Present study | *A.fulgens* |
| H3 | MT891295 | East Sikkim | Sikkim | India | Present study | *A.fulgens* |
| H3 | MT891295 | West Sikkim | Sikkim | India | Present study | *A.fulgens* |
| H3 | MT891295 | East Sikkim | Sikkim | India | Present study | *A.fulgens* |
| H4 | MT891296 | West Sikkim | Sikkim | India | Present study | *A.fulgens* |
| H4 | MT891296 | Neora Valley NP | West Bengal | India | Present study | *A.fulgens* |
| H5 | MT891297 | West Sikkim | Sikkim | India | Present study | *A.fulgens* |
| H5 | MT891297 | Neora Valley NP | West Bengal | India | Present study | *A.fulgens* |
| H5 | MT891297 | Neora Valley NP | West Bengal | India | Present study | *A.fulgens* |
| H5 | MT891297 | East Sikkim | Sikkim | India | Present study | *A.fulgens* |
| H6 | MT891298 | Neora Valley NP | West Bengal | India | Present study | *A.fulgens* |
| H7 | MT891299 | Tawang | Arunachal Pradesh | India | Present study | *A.fulgens* |
| H7 | MT891299 | Tawang | Arunachal Pradesh | India | Present study | *A.fulgens* |
| H7 | MT891299 | Tawang | Arunachal Pradesh | India | Present study | *A.fulgens* |
| H7 | MT891299 | Tawang | Arunachal Pradesh | India | Present study | *A.fulgens* |
| H7 | MT891299 | Mandala | Arunachal Pradesh | India | Present study | *A.fulgens* |
| H7 | MT891299 | Mandala | Arunachal Pradesh | India | Present study | *A.fulgens* |
| H7 | MT891299 | Zingchang, Mandala | Arunachal Pradesh | India | Present study | *A.fulgens* |
| H7 | MT891299 | Fakshee, Lubrang | Arunachal Pradesh | India | Present study | *A.fulgens* |
| H8 | MT891300 | West Sikkim | Sikkim | India | Present study | *A.fulgens* |
| H8 | MT891300 | East Sikkim | Sikkim | India | Present study | *A.fulgens* |
| H8 | MT891300 | Neora Valley NP | West Bengal | India | Present study | *A.fulgens* |
| H9 | MT891307 | Anini, Dibang | Arunachal Pradesh | India | Present study | *A.fulgens* |
| H9 | MT891307 | Anini, Dibang | Arunachal Pradesh | India | Present study | *A.fulgens* |
| H10 | MT891308 | Anini, Dibang | Arunachal Pradesh | India | Present study | *A.fulgens* |
| H10 | MT891308 | Anini, Dibang | Arunachal Pradesh | India | Present study | *A.fulgens* |
| H11 | MT891301 | Zingchang, Mandala | Arunachal Pradesh | India | Present study | *A.fulgens* |
| H12 | MT891302 | Mandala | Arunachal Pradesh | India | Present study | *A.fulgens* |
| H13 | MT891303 | Mandala | Arunachal Pradesh | India | Present study | *A.styani* |
| H14 | MT891304 | Mandala | Arunachal Pradesh | India | Present study | *A.styani* |
| H15 | MT891309 | Lower Dibang | Arunachal Pradesh | India | Present study | *A.styani* |
| H16 | MT891305 | Mechuka, West Siang | Arunachal Pradesh | India | Present study | *A.styani* |
| H17 | MT891306 | Khellong | Arunachal Pradesh | India | Present study | *A.styani* |
| H18 | MT891310 | Anini, Dibang | Arunachal Pradesh | India | Present study | *A.styani* |
| H18 | HQ992979.1 | Gaoligong | Western Yunnan/North Myanmar | China | Hu et al. 2011 | Both |
| H18 | AY849718.1 | Gaoligong | South Yunnan | China | Li et al. 2005 | A.fulgens |
| H19 | HQ992985.1 | South Tibet | Tibet | China | Hu et al. 2011 | A.fulgens |
| H19 | AY849727.1 | South Tibet | Tibet | China | Li et al. 2005 | A.fulgens |
| H20 | HQ992977.1 | Gaoligong | Western Yunnan/North Myanmar | China | Hu et al. 2011 | Both |
| H20 | AY849717.1 | Gaoligong | South Yunnan | China | Li et al. 2005 | A.fulgens |
| H21 | AY849729.1 | South East Tibet | Tibet | China | Li et al. 2005 | A.fulgens |
| H22 | HQ992978.1 | Gaoligong | Western Yunnan/North Myanmar | China | Hu et al. 2011 | Both |
| H23 | HQ992973.1 | Qionglai | North Sichuan | China | Hu et al. 2011 | A.styani |
| H24 | AY849725.1 | Gaoligong | Western Yunnan/North Myanmar | China | Li et al. 2005 | A.fulgens |
| H25 | AF291584.2 | Qionglai | North Sichuan | China | Li et al. 2005 | A.styani |
| H26 | HQ992976.1 | Gaoligong | Western Yunnan/North Myanmar | China | Hu et al. 2011 | Both |
| H26 | AY849732.1 | Gaoligong | Western Yunnan/North Myanmar | China | Li et al. 2005 | A.fulgens |
| H27 | AF291581.2 | Qionglai | North Sichuan | China | Li et al. 2005 | A.styani |
| H28 | HQ992984.1 | South Eastern Tibet | Tibet | China | Hu et al. 2011 | A.fulgens |
| H29 | HQ992982.1 | Gaoligong | Western Yunnan/North Myanmar | China | Hu et al. 2011 | Both |
| H30 | HQ992980.1 | Gaoligong | Western Yunnan/North Myanmar | China | Hu et al. 2011 | Both |
| H31 | HQ992975.1 | Gaoligong | Western Yunnan/North Myanmar | China | Hu et al. 2011 | Both |
| H32 | HQ992974.1 | Xiaoxiangling | South central Sichuan | China | Hu et al. 2011 | A.styani |
| H33 | HQ992970.1 | Liangshan | South Sichuan | China | Hu et al. 2011 | A.styani |
| H34 | HQ992969.1 | Xiaoxiangling | South central Sichuan | China | Hu et al. 2011 | A.styani |
| H35 | HQ992965.1 | Liangshan | South Sichuan | China | Hu et al. 2011 | A.styani |
| H36 | AY849734.1 | Gaoligong | Western Yunnan/North Myanmar | China | Li et al. 2005 | A.fulgens |
| H37 | AY849715.1 | Xiaoxiangling | South central Sichuan | China | Li et al. 2005 | A.styani |
| H38 | AF291583.2 | Qionglai | North Sichuan | China | Li et al. 2005 | A.styani |
| H39 | HQ992983.1 | Gaoligong | Western Yunnan/North Myanmar | China | Hu et al. 2011 | Both |
| H39 | HQ992972.1 | Liangshan | South Sichuan | China | Hu et al. 2011 | A.styani |
| H39 | AY849730.1 | Liangshan | South Sichuan | China | Li et al. 2005 | A.styani |
| H40 | HQ992968.1 | Liangshan | South Sichuan | China | Hu et al. 2011 | A.styani |
| H41 | AY849721.1 | Liangshan | South Sichuan | China | Li et al. 2005 | A.styani |
| H41 | AF291580.2 | Xiaoxiangling | South central Sichuan | China | Li et al. 2005 | A.styani |
| H42 | AY849716.1 | Gaoligong | South Yunnan | China | Li et al. 2005 | A.fulgens |
| H43 | AF291585.2 | Qionglai | North Sichuan | China | Li et al. 2005 | A.styani |
| H44 | AF291579.2 | Liangshan | South Sichuan | China | Li et al. 2005 | A.styani |
| H45 | HQ992967.1 | Qionglai | North Sichuan | China | Hu et al. 2011 | A.styani |
| H45 | AY849719.1 | North Yunnan | Yunnan | China | Li et al. 2005 | A.fulgens |
| H46 | HQ992966.1 | Qionglai | North Sichuan | China | Hu et al. 2011 | A.styani |
| H47 | AY849733.1 | Gaoligong | Western Yunnan/North Myanmar | China | Li et al. 2005 | A.fulgens |
| H48 | AY849731.1 | Gaoligong | Western Yunnan/North Myanmar | China | Li et al. 2005 | A.fulgens |
| H49 | AF291582.2 | Liangshan | South Sichuan | China | Li et al. 2005 | A.styani |
| H50 | HQ992971.1 | Liangshan | South Sichuan | China | Hu et al. 2011 | A.styani |
| H51 | AF291586.2 | Qionglai | North Sichuan | China | Li et al. 2005 | A.styani |
| H52 | HQ992964.1 | Qionglai | North Sichuan | China | Hu et al. 2011 | A.styani |
| H53 | HQ992981.1 | Gaoligong | Western Yunnan/North Myanmar | China | Hu et al. 2011 | Both |
